# Topological features of gene regulatory networks predict patterns of natural diversity in environmental response

**DOI:** 10.1101/080804

**Authors:** David L. Des Marais, Rafael F. Guerrero, Jesse R. Lasky, Samuel V. Scarpino

## Abstract

Molecular interactions affect the evolution of complex traits. For instance, adaptation may be constrained by pleiotropic or epistatic effects, both of which will be reflected in the structure of molecular interaction networks. To date, empirical studies investigating the role of molecular interactions in phenotypic evolution have been idiosyncratic, offering no clear patterns. Here, we investigated the network topology of genes putatively involved in local adaptation to two abiotic stressors—drought and cold—in *Arabidopsis thaliana*. Our findings suggest that the gene-interaction topologies for both cold and drought stress response are non-random, with genes that show genetic variation in drought response (GxE) being significantly more peripheral and cold response genes being significantly more central than genes not involved in either response. We suggest that the observed topologies reflect different constraints on the genetic pathways involved in the assayed phenotypes. The approach presented here may inform predictive models linking genetic variation in molecular signaling networks with phenotypic variation, specifically traits involved in environmental response.

**Significance Statement:** Our study focuses on genes whose transcriptional activity exhibits genetic variation in response to the environment, or “GxE.” GxE is a widely observed phenomenon of critical importance to understanding the genotype-to-phenotype map, the evolution of natural populations, medical genetics, population response to climate change, and agricultural improvement. We investigated expression GxE in plant responses to two abiotic cues: cold and drought. We found that genes showing genetically variable response to cold stress are centrally located in regulatory networks whereas genes showing genetically variable response to drought stress are peripherally located in regulatory networks. This result suggests that selection is presented with vastly different mutational landscapes for shaping evolutionary or breeding response to these two important climatic factors

## Introduction

Genes do not function nor evolve in isolation. The transcriptional activities of genes in a genome are often highly correlated with one another, forming hierarchical regulatory networks comprised of functionally related modules (1). Within such networks, some genes – “nodes” - have stronger or more interactions – “edges” - with one another than do other genes. Because the effect size of mutations is strongly associated with their evolutionary fate (2), the structural properties of genetic regulatory networks will likely affect selection acting on individual component genes (3-5). Advances in high-throughput molecular phenotyping and systems analysis have improved our ability to characterize molecular interaction networks, providing the opportunity to address classic questions about the evolution of genetic interactions.

Two related features of gene regulatory networks might affect the evolution of individual genes within those networks. The first is the widespread observation that genes vary in their number of interacting neighbor genes, perhaps even by orders of magnitude (6). This feature – the centrality or connectivity of a gene – can be measured in many different ways, including the number of directly interacting genes or the number of paths to other genes that pass through a given gene (7). The second feature of networks that can impact gene evolution is modularity, i.e. the degree to which the network is composed of functionally related sub-networks of genes, or modules. Modules are often under the transcriptional control of core proteins, high-level switches that regulate the module’s activity (8) using shared regulatory motifs among genes within it (9). Evidence for the pleiotropic nature of core genes has been found through decades of developmental genetics research, which identified putative master regulators of the level, timing, and location of expression of tens to thousands of other genes (3, 8, 10).

Transcriptional regulation by core genes plays an important role in adaptive responses to the environment (11, 12). Environmentally responsive transcripts are often co-regulated as functional modules (13, 14), and in some instances the response of particular genes to environmental cues may be characteristic of entire species or kingdoms (15, 16). Considerable genetic variation in transcriptional response to environment – expression Genotype by Environment interaction, eGxE – has also been identified within species (17). At the molecular level, eGxE may be controlled by genetic variants acting in cis, e.g. by SNP or presence-absence variants in promoter motifs, or by genetic variants acting in trans, such as transcription factors, small RNA species, or a number of other regulatory factors upstream of genes showing eGxE. Genetic variants affecting eGxE are of particular interest because GxE represents the mutational substrate for the evolution environmental response (18, 19) and because GxE for fitness is required for local adaptation (20).

In the present study, we explore two hypotheses for how environmentally responsive regulatory networks evolve and might thereby be involved in local adaptation to environment. The first hypothesis, that eGxE is driven by genetic variation in core transcriptional regulatory proteins, arises from the observation that suites of traits often show high genetic correlation ((21); in this context, “traits” could be either individual transcripts or higher-level physiological or developmental phenotypes). Genetic variants in one or a small number of regulatory genes could therefore have considerable downstream consequences, both positive and negative with respect to transcription level, trait expression, and fitness. This model predicts that eGxE genes would have relatively high network connectivity and, by extension, be clustered in relatively discrete functional modules. The second hypothesis posits that eGxE could be primarily driven by variation in genes located peripherally in transcriptional networks, which are expected to have smaller effect sizes and reduced deleterious pleiotropy. Variation in peripheral genes could therefore allow natural selection to “fine-tune” environmental response by changing only a small number of expression or higher-order traits. While these are not mutually exclusive hypotheses, their relative importance in nature has not been established.

Here, we extend earlier work assessing genetic variation in transcriptional activity during acclimation to cold (22) and soil drying (23) in *Arabidopsis thaliana*. We predict different patterns for the genes associated with each environmental response based on our previous observations. The sequence conservation in environmentally responsive promoter motifs among cold eGxE genes in Arabidopsis (24) leads us to predict that cold acclimation eGxE genes will be clustered and highly connected. The conserved patterns of nucleotide diversity imply that *cis*-regulatory elements of eGxE genes for cold are evolutionarily conserved and that expression diversity is controlled by genetic diversity in upstream regulators such as transcription factors. These transcription factors, acting as core regulators, would cause the coexpression of large sets of eGxE genes that would be observed as a highly connected network. Conversely, we predict that drought eGxE genes will be located peripherally in networks because these genes show evidence of adaptive *cis*-regulatory variation (24). This predominant role of *cis* variants suggests that most genes involved in drought local adaptation have small *trans* effects and are therefore likely to have low connectivities in the coexpression network. By extension, we predict that the relative contribution of cis- and trans- associated expression variation indicates the structure of response to environment across a molecular network.

## Results and Discussion

### Stress-responsive genes are non-randomly distributed in a transcriptional co-expression network

We first tested the hypothesis that genes whose expression responds to environmental gradients are non-randomly distributed in a transcriptional regulatory network. Specifically, we assessed the positions of stress-responsive genes in a co-expression network estimated for *A. thaliana* by Feltus et al. ((25); hereafter “Feltus network”). We take stress-responsive genes from a previous analyses of two experimental datasets: our “cold” dataset comes from Lasky et al ((24) a re-analysis of Hannah et al. (22)), while our “drought” dataset comes from Des Marais et al. (23). Briefly, the approach taken by Lasky et al. (24) and Des Marais et al. (23) was to partition variance in gene expression level among the effects of genotype (inbred natural accession), environment (response to experimental treatment), and their interaction (“eGxE”). In the context of this paper, stress-responsive genes are those classified as “eGxE”, which show a genetically variable response to environmental changes, as well as genes that show a similar pattern of response across natural accessions “cold-response” or “drought-response” genes).

The Feltus network was reconstructed by aggregating data from 7105 published microarray experiments differing in environment, tissue, and genotype; this network hypothesis is therefore a meaningful summary of the transcriptional relationships among genes in diverse environmental settings and genomic backgrounds. For each gene node) in the network, we calculated the degree of the node, which measures the number of neighboring nodes in the network, as well as the centrality if the node. Centrality of a node estimates the number of paths between other nodes that pass through this node; here, we present node centrality as eigenvector centrality. A gene will have a high eigenvector centrality if it is both well connected itself and if its neighbors are also well connected (see Methods and Newman (26) for more details). We obtained qualitatively similar results with alternative centrality metrics, e.g., degree, betweeness, and closeness centrality (see Supplement).

For both data sets, genes showing significant eGxE were non-randomly distributed with respect to network degree and centrality. Drought eGxE genes had lower degrees (Figure 1a) [median for drought: eGxE = 4; non- eGxE = 13], i.e. had fewer connections to other genes, and were less central (Figure 1b) when compared to non- eGxE genomic controls. Cold eGxE genes exhibited the opposite effect, having higher degree (Figure 1c) [median cold: eGxE = 38; non- eGxE = 11] and being more centrally located (Figure 1d) compared to genomic controls. Statistical significance was determined by selecting a random subset of genes equal in size to the genes showing eGxE and then calculating their degree and centrality (see Supplement). Out of 10,000 permutations, we did not observe a single set of genes with more extreme low (drought) or high (cold) distributions of degree and eigenvector centrality, corresponding to a p-value of 10^−4^. Interestingly, cold eGxE genes had higher average connectivity than that of cold or drought response (environment) genes, of which we have hypothesized represent the conserved environmental response of a species (23, 24) (see Supplement).

**Figure 1.**
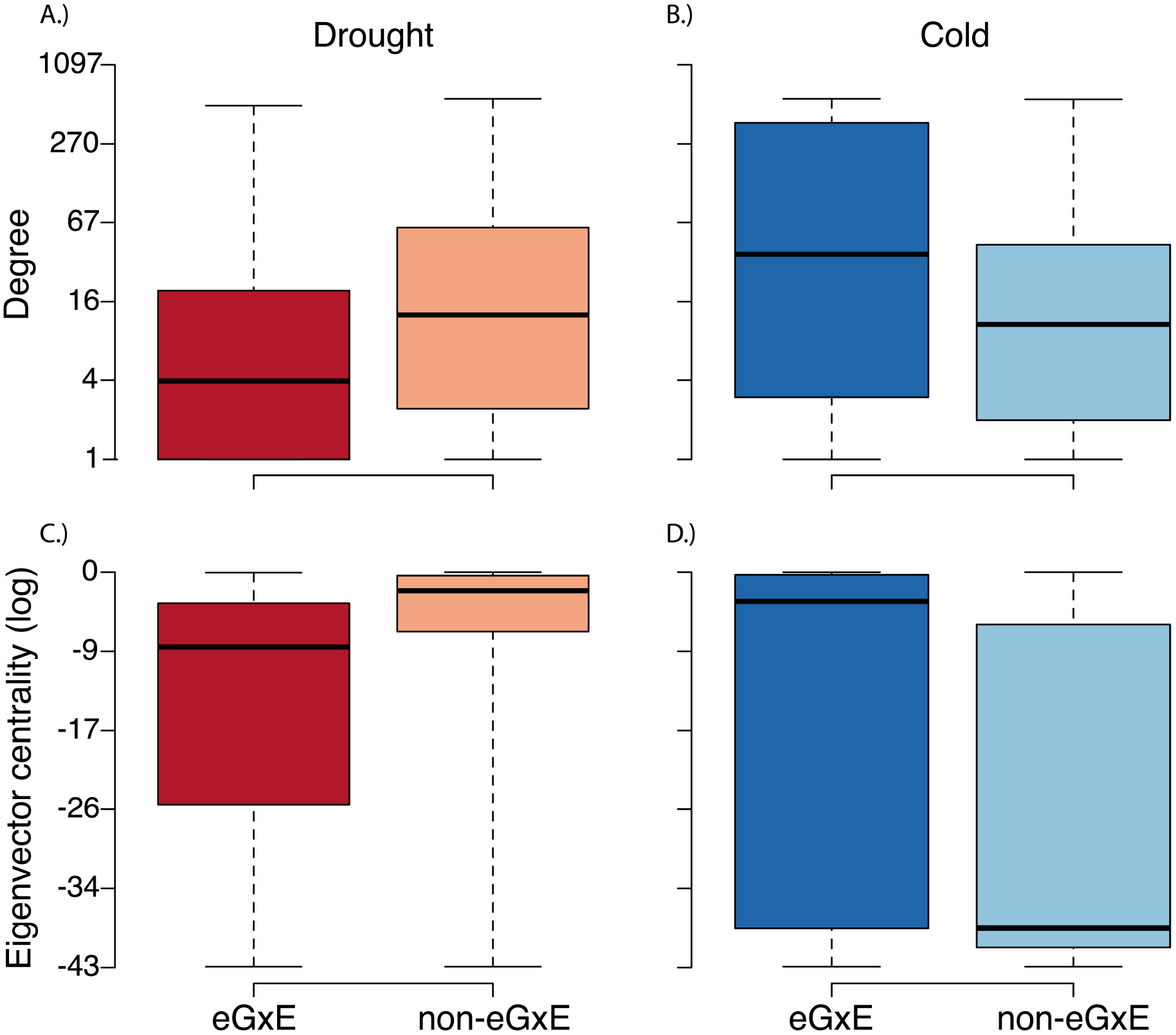
Expression GxE genes are non-randomly distributed in the Arabidopsis gene regulatory network. As a group, eGxE genes in drought conditions have significantly lower degree A) and eigenvector centrality C) than do genomic controls. As a group, eGxE genes in cold condition have significantly higher degree (B) and eigenvector centrality (D) than do genomic controls. The solid line represents the median, colored boxes indicate the interquartile range, and whiskers mark the entire range of the data.

To verify the robustness of our results, we confirmed the non-random distribution of genes using community detection. The above analysis using the Feltus network makes the assumption that all genes in the network are equally likely to participate in environmental response. However, because certain pathways *must* be involved in the phenotypic response to drought or cold, it is possible that the appropriate null distribution should be constructed using nearby genes e.g. those with strong co-expression in the Feltus network). To explore this possible statistical artifact, we performed standard community detection using the leading eigenvector method (27) as implemented in the R package igraph v.1.0.1 (28) on the Feltus network and subdivided the graph into sets of genes that are densely connected among themselves and loosely connected to other parts of the gene co-expression network. Averaging the results across all detected communities, with significance again determined by permutation test, we recover the same pattern seen in the global network: Cold eGxE genes have higher degree cold eGxE genes had a median of 34 more connections than non eGxE genes) and a significantly higher median eigenvector centrality (median 0.14 above non eGxE), while drought eGxE genes have lower degree (cold eGxE genes had a median of 16 fewer connections than non eGxE genes) and had a significantly lower eigenvector centrality median 0.04 below non eGxE). A similar result was found using the subnetworks (their “GILs”) previously constructed by Feltus see Supplement).

We further validated this result by performing iterative out-of-sample model validation. Briefly, we randomly selected 80% of genes in the network and constructed a generalized linear model with a binomial error distribution i.e. a logistic regression) to predict genes as eGxE based solely on their degree and eigenvector centrality. We then predicted the eGxE state for the remaining 20% of genes and recorded the error. We repeated this procedure 1,000 times for both cold and drought. Assuming a threshold for accurate classification of 5%, we were able to correctly classify 95.4% of genes for cold and 77.0% of genes for drought. These results accommodate classification errors for both eGxE and non- eGxE genes, which means that for cold we were able to correctly classify nearly every gene included in the co-expression network as being eGxE based solely on its degree and eigenvector centrality.

### eGxE genes show modular distribution that differs between environments

We next asked whether the non-random distribution of node connectivity of eGxE genes reflects their membership in particular sub-communities, or modules, of interacting genes as identified using our community detection approach, described above. Interestingly, both the cold and drought eGxE genes were non-randomly distributed with respect to the sub-communities defined using our community detection approach. For cold, 32.5% of all eGxE genes exist within a single, large sub-community containing 605 genes (Figure 2a) and an additional 26.5% of cold eGxE genes are found in a second large sub-community containing 425 genes. In contrast, for drought, the two sub-communities with highest accumulation of eGxE genes together contain only 18% of the total number of eGxE genes (Figure 2b shows the larger of these two). Moreover, drought eGxE are statistically over-represented in five small sub-communities comprised of between 10 and 100 members (Figure 3b), while cold eGxE genes are clustered in a few large communities (Figure 3a). The membership of eGxE genes in sub-communities of differing size recapitulates our earlier result: cold eGxE genes tend to be functionally connected to many other genes, while genes involved in drought response tend to be in peripheral network positions.

**Figure 2.**
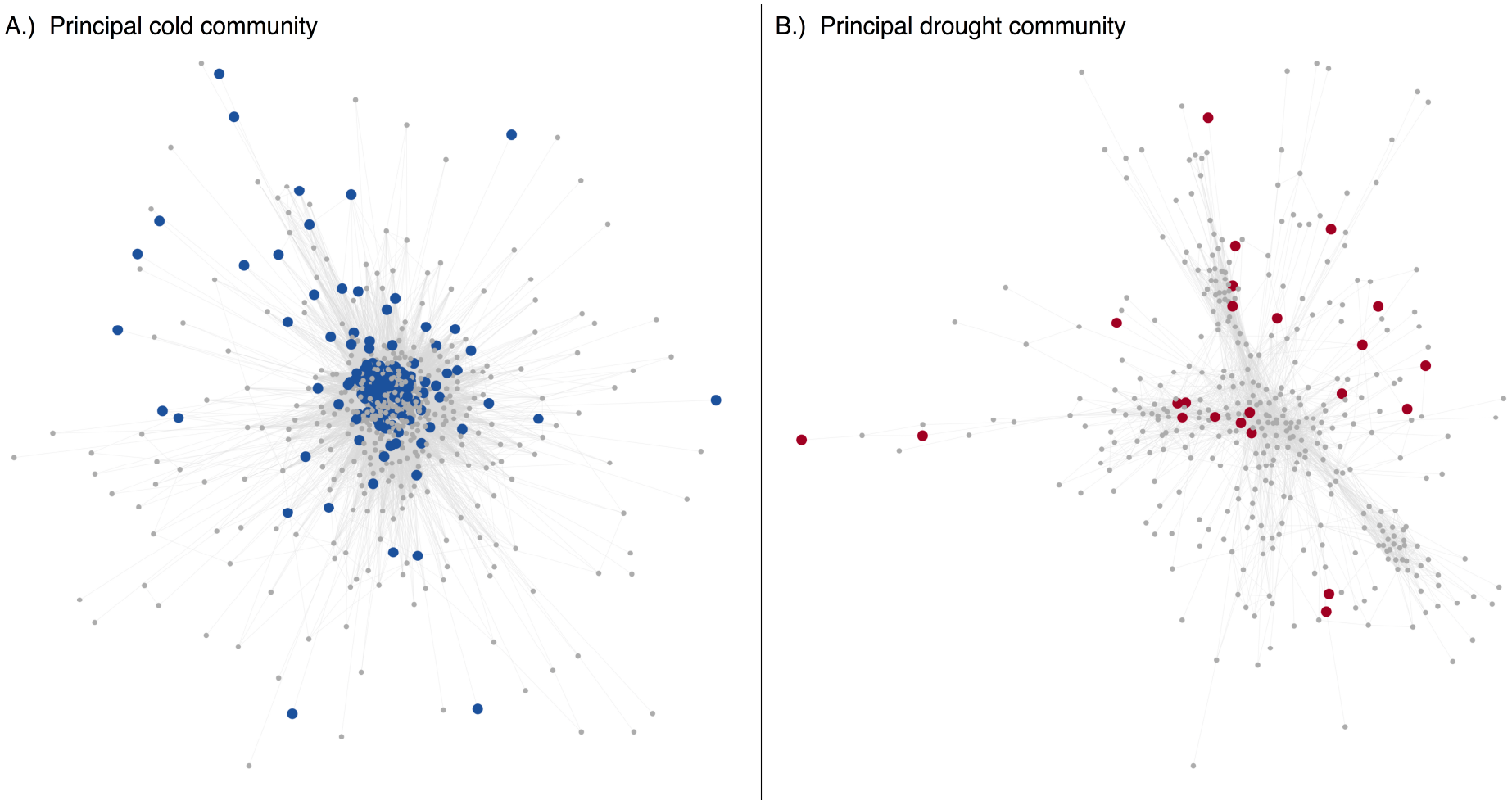
Graphical representation of two sub-communities of the Arabidopsis gene regulatory network showing the sub-community with the highest over-representation of cold eGxE genes (A) and the sub-community with the highest over-representation of drought eGxE genes B). Colored circles indicate genes showing significant eGxE at pFDR=0.05 from (23, 24).

**Figure 3.**
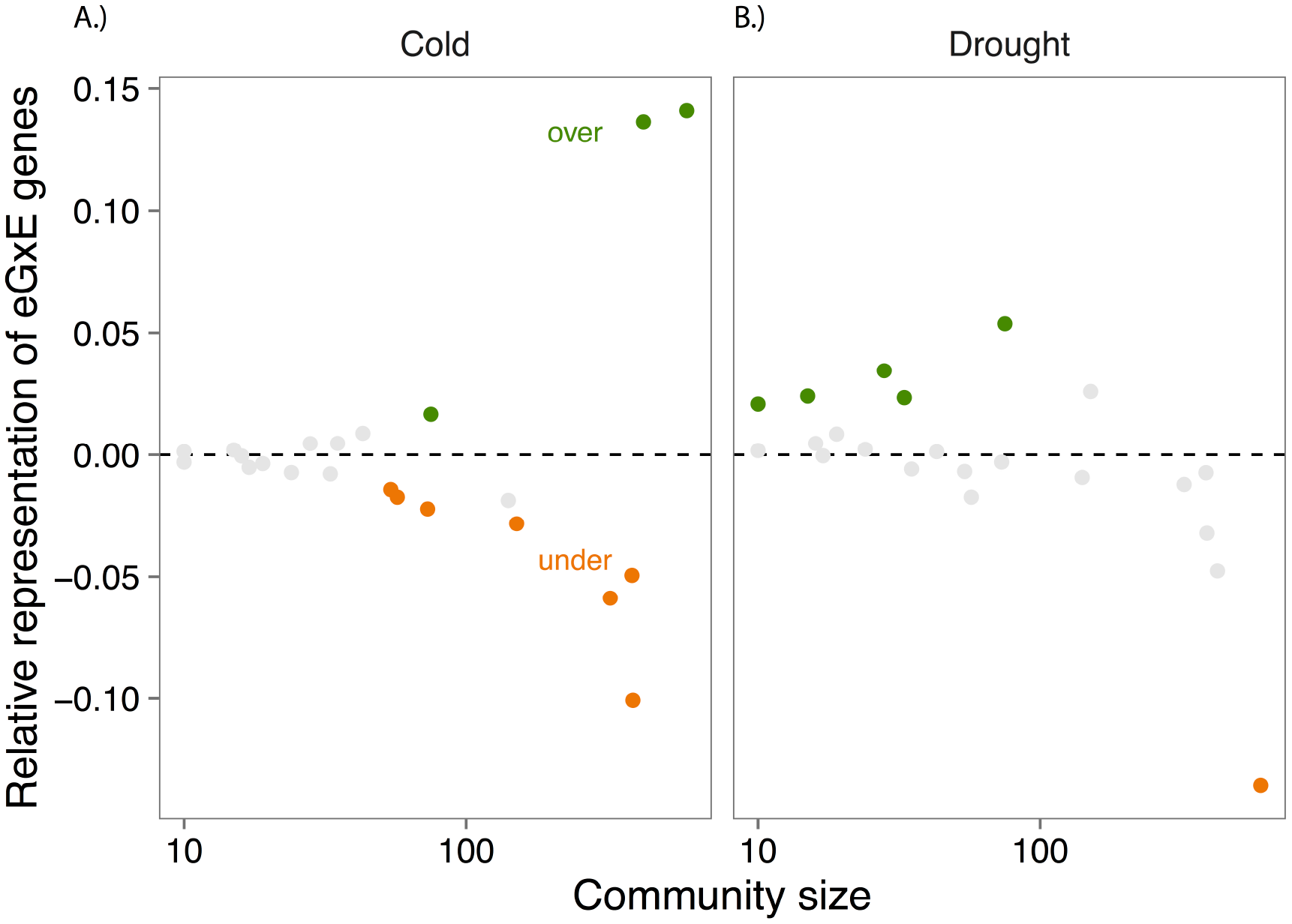
Cold A) and drought B) eGxE genes have different distributions across sub-communities of the gene regulatory network. Cold eGxE genes are overrepresented in two large sub-communities, each containing over 400 total genes. Five smaller sub-communities (<100 genes) are enriched for drought eGxE genes. The vertical axis is the difference between the observed and expected number of eGXE genes in the community, relative to the total number of genes in the community. Each sub-community in the analysis is shown with a circle only subcommunities with ten or more genes are included). Green circles indicate sub-communities in which eGxE genes are significantly overrepresented, while orange circles represent subcommunities with significant underrepresentation of eGxE genes (Fisher’s exact test; FDR=0.05).

To test the hypothesis that the sub-communities with diverging patterns of expression eGxE reflect natural variation in function, we took the genes in the two sub-communities with the most overrepresentation in cold and in drought response and tested for enrichment of gene ontology (GO) annotations. We found 116 significant GO terms enriched in the most over-represented cold eGxE sub-community. The top seven terms were all related to photosynthesis and related processes, and the next two terms were for response to abiotic stimulus and response to cold (Table S3). Altered primary metabolism is frequently observed during cold acclimation, in part due to the accumulation of sugars as cryoprotectants (29, 30), so this eGxE may reflect that some of the sampled accessions modify metabolism during cold response to a different degree than do other accessions. We found 89 significant GO terms enriched in the most over-represented drought eGxE subcommunity. The top term and many of the subsequent terms were for immune and defense responses (Table S4; many genes, particularly kinases, annotated as immune and defense responses also show responses to abiotic stress (31)). Previously, we found that drought eGxE genes showed very few significant functional enrichments using a genome-wide test for statistical enrichment (23), suggesting that the network-informed approach used here may afford additional statistical power to detect functional patterns in these high-dimensional datasets.

### The evolution of gene expression response to the environment

Previously, we demonstrated an important role of cis-regulatory variants underlying diversity of environmental response among natural genotypes of Arabidopsis (24). Natural variation in response to drought showed different genomic patterns than did natural variation in response to cold, suggesting that natural selection may affect different parts of the transcriptional regulatory networks for these two complex traits. Specifically, the proximal promoters of genes showing eGxE for drought had significantly higher nucleotide diversity and significantly higher among-genotype variation in key drought-responsive promoter motifs abscisic acid responsive elements, ABREs) when compared to genome averages (24). These earlier observations for eGxE drought genes are consistent with the results presented here: drought eGxE genes are in smaller modules and are relatively lowly connected to other genes, suggesting that genetic variation in expression response to the environment is controlled locally, possibly by a large number of cis-acting variants. This architecture may permit functionally diverse modules to act independently from one another, i.e. showing environmental response in only some genotypes (32). Our results may also explain why expression QTL (eQTL) studies of drought response identify a preponderance of cis- acting eQTL and few trans- eQTL (33, 34).

Collectively, these observations suggest two, non-exclusive, hypotheses for how natural populations of plants adapt to variation in soil water content. First, diverse populations and species acclimate to transient soil drying stress in diverse ways – via changes in growth, transpiration, leaf area-volume ratios, timing of reproduction, cell wall composition, and synthesis of various osmoprotectants and chaperonins, to name but a few (35). In this model, the modular regulatory architecture observed here reflects functionally different transcriptional modules driving these different physiological responses. The extent to which such physiological alterations are under independent or common genetic control is presently unknown. Second, because most plants likely experience fluctuations in water availability on a daily or seasonal basis (36-38), plants must maintain some degree of physiological response to drying. Conditionally-active regulatory alleles may therefore allow for fine-tuning of response because they could provide fitness benefits in one environment without deleterious consequences in second environment “yield-drag”; (39)). Indeed, cis-acting eGxE variants for drought response are predominantly variance-changing (i.e. conditionally neutral or differentially sensitive to conditions) rather than direction-changing (33). Moreover, the vast majority of QTL showing GxE in studies of abiotic stress are also conditionally neutral, or show differential sensitivity to environmental cues (17).

Nucleotide diversity in the proximal promoters of cold eGxE genes is also elevated but, in contrast to drought genes, not to a degree that is statistically significant compared to genome averages (24). The promoters of cold eGxE genes exhibit lower among-genotype turnover of known cold-responsive motifs compared to genome averages (specifically, the c-repeat binding factor/dehydration responsive elements, CRT/DREs). Along with our observation that cold eGxE genes are highly connected in the transcriptional regulatory network, these patterns of sequence diversity suggest that the transcriptional control of eGxE for cold acclimation is driven by genetic variants in upstream regulatory features, such as transcription factors. Natural variants in transcription factors are expected to have larger mutational effect size due to the regulatory influence of these proteins on downstream genes.

It is currently unknown what drives the apparent difference in the genetic architecture of natural variation in response to drought compared to response to cold. While daily or seasonal drying stress is likely experienced by *A. thaliana* plants to some degree across the species range, severe cold stress is likely only experienced by populations at higher latitudes or altitudes. A recent study demonstrated that multiple, apparently independent, loss of function mutations in key transcriptional regulators of cold-responsive genes are associated with geographic variation in winter temperature across the range of *A. thaliana* (40). The activity of these CBF transcription factors shows a strong positive correlation with the capacity of *A. thaliana* natural accessions to acclimate to cold (22, 41, 42). Perhaps cold-associated selective gradients involve sharp transitions (along spatial and climatic gradients) in optimal phenotypes, with only a few fitness optima for cold tolerance - manifested as cold-tolerant, functionally CBF versus cold-intolerant, non-functional CBF accessions. By contrast, drought tolerance may be subject to smoother, more gradual selective gradients, for which many smaller-effect variants allow fine-tuning of response.

### Conclusions

Our results suggest that topological relationships among genes in transcriptional regulatory networks affect how natural populations adapt to the multivariate environment. A promising extension of our approach is to link information regarding the topological features of a given gene - its connectivity, in the case presented here, as well as its membership in particular functional modules - with information associating genetic variants with phenotype from genetic mapping. Such a combined analysis could clarify how putatively functional variants identified via association mapping result in phenotypic variation via cellular and physiological mechanisms (43).

From an applied perspective, understanding how natural variation affects transcriptional regulatory networks may inform decisions about how to improve agricultural performance in challenging environments. We note that breeding for improved performance under soil drying has been quite challenging (44); our results suggest that targeting specific physiological mechanisms by manipulating genes at the “tips” of regulatory networks, shown herein to exhibit drought eGxE, may be a more fruitful strategy than targeting central regulatory molecules which may exhibit undesirable pleiotropic effects “yield drag;” e.g. (45)).

## Methods

### Co-expression and regulatory networks

We used two published datasets on gene expression interactions in in Arabidopsis thaliana. The first dataset was global i.e. genome-wide and not restricted to certain functions or pathways). We used the global co-expression network and 86 subcomponent genome-wide gene coexpression networks created by (25). The authors first obtained 7,105 publicly available ATH1 Affymetrix microarray samples and applied a thresholding algorithm (random matrix theory) to generate a global network containing 3,297 nodes and 129,134 edges. These nodes represent 16% of Arabidopsis genes on the ATH1.

A complication from this approach arises because of interactions between genotype, expression networks and environment including ontogeny, tissue, or cell type; (46)). Thus a prior step of partitioning expression data may help to account for some of this heterogeneity and better reveal co-expression networks. Through k-means partitioning, Feltus et al. (25) generated 86 gene interaction layers (GILs), i.e. 86 smaller, non-global networks, that together included 19,588 genes, which represents 95% of Arabidopsis genes on the ATH1.

### Data: transcriptomic responses to cold and drought stress

We used two published studies on natural variation in transcriptomic response to cold (22) and drought (23). Both studies used the ATH1 microarray to estimate genome-wide transcript abundance. Each study subjected a diverse panel of 9 (22) or 17 (23) natural accessions to a cold or drought treatment, respectively. Lasky et al. (2014) re-analyzed the Hannah et al. dataset to match the analyses by Des Marais et al. In brief, those authors modeled transcript abundance using factorial ANOVA including Accession (i.e. Genotype), Treatment, and their interaction and identified significantly differentially expressed gene models at pFDR of 0.05.

### Network Methods

We quantified the degree to which a gene was central using four standard network metrics, 1.) degree (raw number of connections), 2.) closeness centrality (the inverse of the average shortest path between the focal gene and all other genes in the network), 3.) betweenness centrality (number of shortest paths between all pairs of nodes in the network, which pass through the focal node), and 4.) eigenvector centrality. Eigenvector centrality, which is closely related to Google’s PageRank algorithm (47) measures a node’s centrality based on both the node’s own position in the network and the position of that node’s neighbors in the network. More specifically, a node’s eigenvector centrality will be proportional to the average centralities of its neighbors (26)

To identify community structure and assign genes to communities, we used the leading eigenvalue algorithm. Our goal was to determine whether there are groups of genes, which are more connected to each other than they are to other genes, referred to as community detection (48). A variety of methods exist for performing community detection, but we selected the leading eigenvalue approach because it is computationally efficient on large networks. Briefly, the adjacency matrix of the network is corrected based on the expected number of edges in a random graph, using the configuration model, then the distribution of eigenvalues and the loading of nodes onto eigenvectors can be used to 1.) determine whether evidence exists for the presence of modular communities and 2.) assuming such structure exists, assign genes to communities, see Newman 2006 for more details. All of the analyses, i.e. calculation of centrality measures and community detection, was performed using the R package igraph v.1.0.1 (28).

### Enrichment analyses

We tested whether genes with multiple types of expression response to abiotic stress exhibited non-random network metrics. We also tested whether genes exhibiting high Fst exhibited non-random network metrics. Determining whether a gene has a higher value for any of these metrics is not appropriate for parametric stats. Therefore, we conducted permutations to generate null expectations. The natural accessions used in this study all show varying degrees of sequence divergence and gene gain/loss as compared to the reference Col-0 genome, which was used to generate the microarrays used in these experiments. This variance could generate spurious “gene-by-environment interaction” for gene expression. We therefore used a strict filtering scheme to exclude genes that had polymorphisms in ATH1 probe sites (23). In order to assess the biological function of regulatory communities, we first identified communities containing the greatest proportion of GxE genes for each abiotic stressor. We then tested gene ontology (GO) term enrichment in each of these communities (AgriGO).

## Acknowledgements

We thank R. Hopkins and E. Josephs for comments on the manuscript. T. Juenger and J. G. Monroe provided stimulating discussion on the genetic architecture of abiotic stress response and expression GxE. This work is a product of the “Molecular Networks and Evolution Across Biological Scales” workshop supported by the Santa Fe Institute.

## References

1. Wagner GP & Zhang J (2011) The pleiotropic structure of the genotype-phenotype map: the evolvability of complex organisms. Nature Reviews Genetics 12 (3):204–213.

2. Fisher RA (1930) The Genetical Theory of Natural Selection (Clarendon Press, Oxford).

3. Erwin DH & Davidson EH (2009) The evolution of hierarchical gene regulatory networks. Nature Reviews Genetics 10:141–148.

4. Jeong H, Mason SP, Barabasi AL, & Oltvai ZN (2001) Lethality and centrality in protein networks. Nature 411:41.

5. Rausher MD, Miller RE, & Tiffin P (1999) Patterns of Evolutionary Rate Variation Among Genes of the Anthocyanin Biosynthetic Pathway. Molecular Biology and Evolution 16 (2):266–274.

6. Guelzim N, Bottani S, Bourgine P, & Kepes F (2002) Topological and causal structure of the yeast transcriptional regulatory network. Nat Genet 31 (1):60–63.

7. Lin W-h, Liu W-c, & Hwang M-j (2009) Topological and organizational properties of the products of house-keeping and tissue-specific genes in protein-protein interaction networks. BMC systems biology 3:32.

8. Wilkins AS (2002) The evolution of developmental pathways Sinauer, Sunderland, MA).

9. Wray GA, et al. (2003) The evolution of transcriptional regulation in eukaryotes. Molecular Biology and Evolution 20 (9)

10. Wagner GP, Pavlicev M, & Cheverud JM (2007) The road to modularity. Nat Rev Genet 8 (12):921–931.

11. Gibson G (2008) The environmental contribution to gene expression profiles. Nature Reviews Genetics 9:575–581.

12. Weake VM & Workman JL (2010) Inducible gene expression: diverse regulatory mechanisms. Nature Reviews Genetics 11:426–437.

13. Fowler S, Cook D, & Thomashow MF (2007) The CBF cold-response pathway. Plant Abiotic Stress, eds Jenks MA & Hasegawa PM (Wiley-Blackwell, Chichester), pp 7199.

14. Luscombe NM, et al. (2004) Genomic analysis of regulatory network dynamics reveals large topological changes. Nature 431:308–312.

15. McCarroll SA, et al. (2004) Comparing genomic expression patterns across species identifies shared transcriptional profile in aging. Nat Genet 36 (2):197–204.

16. Carvallo MA, et al. (2011) A comparison of the low temperature transcriptomes and CBF regulons of three plant species that differ in freezing tolerance: *Solanum commersonii, Solanum tuberosum*, and Arabidopsis thaliana. J Exp Bot 62 (11):3807–3819.

17. Des Marais DL, Hernandez KH, & Juenger TE (2013) Genotype-by-environment interaction and plasticity: exploring genomic responses of plants to the abiotic environment. Annual Review of Ecology, Evolution and Systematics 44:5–29.

18. Schlichting CD & Pigliucci M (1998) Phenotypic Evolution: A Reaction Norm Perspective Sinauer Associates, Sunderland, MA).

19. Via S & Lande R (1985) Genotype-environment interaction and the evolution of phenotypic plasticity. Evolution 39:505–523.

20. Kawecki TJ & Ebert D (2004) Conceptual issues in local adaptation. Ecology Letters 7:1225–1241.

21. Lynch M & Walsh B (1998) Genetics and Analysis of Quantitative Traits (Sinauer, Sunderland, MA).

22. Hannah MA, et al. (2006) Natural genetic variation of freezing tolerance in Arabidopsis. Plant Physiol 142 (1):98–112.

23. Des Marais DL, et al. (2012) Physiological genomics of response to soil drying in diverse Arabidopsis accessions. Plant Cell 24 (3):893–914.

24. Lasky JR et al. (2014) Natural variation in abiotic stress responsive gene expression and local adaptation to climate in Arabidopsis thaliana. Molecular Biology and Evolution 31:2283–2296.

25. Feltus FA, Ficklin SP, Gibson SM, & Smith MC (2013) Maximizing capture of gene co-expression relationships through pre-clustering of input expression samples: An *Arabidopsis* case study. BMC systems biology 7:44.

26. Newman ME (2008) The mathematics of networks. The new palgrave encyclopedia of economics 2 (2008):1–12.

27. Newman ME (2006) Finding community structure in networks using the eigenvectors of matrices. Phys Rev E Stat Nonlin Soft Matter Phys 74 (3 Pt 2):036104.

28. Csardi G & Nepusz T (2006) The igraph software package for complex network research. International Journal of Complex Systems 1695 (5):1–9.

29. Thomashow M (1999) Plant cold acclimation: freezing tolerance genes and regulatory mechanisms. Annual Reviews of Plant Physiology and Plant Molecular Biology 50:571–599.

30. Colton-Gagnon K, et al. (2014) Comparative analysis of the cold acclimation and freezing tolerance capacities of seven diploid *Brachypodium distachyon* accessions. Ann Bot 113 (4):681–693.

31. Marshall A, et al. (2012) Tackling drought stress: receptor-like kinases present new approaches. Plant Cell 24(6):2262–2278.

32. Draghi JA, Parsons TL, Wagner GP, & Plotkin JB (2010) Mutational robustness can facilitate adaptation. Nature 463 (7279):353–355.

33. Cubillos FA, et al. (2014) Extensive cis-regulatory variation robust to environmental perturbation in Arabidopsis. Plant Cell 26(11):4298–4310.

34. Lowry DB, et al. (2013) Expression quantitative trait locus mapping across water availability environments reveals contrasting associations with genomic features in Arabidopsis. Plant Cell 25 (9):3266–3279.

35. Chaves MM, Maroco JP, & Pereira J (2003) Understanding plant responses to drought – from genes to the whole plant. Functional Plant Biology 30:239–264.

36. Richards CL, Rosas U, Banta J, Bhambhra N, & Purugganan MD (2012) Genome-wide patterns of Arabidopsis gene expression in nature. PLoS Genet 8 (4):e1002662.

37. Chaves MM, et al. (2002) How Plants Cope with Water Stress in the Field? Photosynthesis and Growth. Annals of Botany 89 (7):907–916.

38. Plessis A, et al. (2015) Multiple abiotic stimuli are integrated in the regulation of rice gene expression under field conditions. Elife 4.

39. Mickelbart MV, Hasegawa PM, & Bailey-Serres J (2015) Genetic mechanisms of abiotic stress tolerance that translate to crop yield stability. Nat Rev Genet 16 (4):237–251.

40. Monroe JG, et al. (2016) Adaptation to warmer climates by parallel functional evolution of CBF genes in Arabidopsis thaliana. Mol Ecol 25 (15):3632–3644.

41. Gehan MA, et al. (2015) Natural variation in the C-repeat binding factor cold response pathway correlates with local adaptation of Arabidopsis ecotypes. Plant J 84 (4):682–693.

42. Gery C, et al. (2011) Natural variation in the freezing tolerance of *Arabidopsis thaliana:* effects of RNAi-induced CBF depletion and QTL localisation vary among accessions. Plant Sci 180 (1):12–23.

43. Rockman MV (2008) Reverse engineering the genotype-phenotype map with natural genetic variation. Nature 456:738–744.

44. Passioura J (2010) Scaling up: the essence of effective agricultural research. Functional Plant Biology 37:585–591.

45. Kasuga M, Liu Q, Miura S, Yamaguchi-Shinozaki K, & Shinozaki K (1999) Improving plant drought, salt and freezing tolerance by gene transfer of a single stress-inducible transcription factor. Nature Biotechnology 17:287–291.

46. Fukushima A, et al. (2012) Exploring tomato gene functions based on coexpression modules using graph clustering and differential coexpression approaches. Plant Physiol 158 (4):1487–1502.

47. Langville AN & Meyer CD (2011) Google’s PageRank and beyond: The science of search engine rankings. (Princeton University Press, Princeton, NJ).

48. Fortunato S & Hric D (2016) Community detection in networks: A user guide. arXiv:1608.00163.

